# Sensorimotor conflicts induce somatic passivity and louden quiet voices in healthy listeners

**DOI:** 10.1101/2020.03.26.005843

**Authors:** Pavo Orepic, Giulio Rognini, Oliver Alan Kannape, Nathan Faivre, Olaf Blanke

**Author notes:** equal contribution (Nathan Faivre, Olaf Blanke). Corresponding author: Olaf Blanke, Bertarelli Chair in Cognitive Neuroprosthetics, Center for Neuroprosthetics & Brain Mind Institute, School of Life Sciences, Campus Biotech, Swiss Federal Institute of Technology, Ecole Polytechnique Fédérale de Lausanne (EPFL), CH – 1012 Geneva.

## Abstract

Sensorimotor conflicts are known to alter the perception of accompanying sensory signals and deficits in sensory attenuation have been observed in schizophrenia. In the auditory domain, self-generated tones or voices (compared to tones or voices presented passively or with sensorimotor delays) have been associated with changes in loudness perception and attenuated neural responses. It has been argued that for sensory signals to be attenuated, predicted and sensory consequences must have a consistent spatiotemporal relationship, between button presses and reafferent tactile signals, via predictive sensory signaling, a process altered in schizophrenia. Here, we investigated auditory sensory attenuation for a series of morphed voices while healthy participants applied sensorimotor stimulations that had no spatiotemporal relationship to the voice stimuli and that have been shown to induce mild psychosis-like phenomena. In two independent groups of participants, we report a loudening of silent voices and found this effect only during maximal sensorimotor conflicts (versus several control conditions). Importantly, conflicting sensorimotor stimulation also induced a mild psychosis-like state in the form of somatic passivity and participants who experienced stronger passivity lacked the sensorimotor loudening effect. We argue that this conflict-related sensorimotor loudness amplification may represent a reduction of auditory self-attenuation that is lacking in participants experiencing a concomitant mild psychosis-like state. We interpret our results within the framework of the comparator model of sensorimotor control, and discuss the implications of our findings regarding passivity experiences and hallucinations in schizophrenia.

## Introduction

Our capacity to process motor signals, their reafferent sensory consequences, and sensory prediction signals is crucial for motor control and perception^1^ and for updating internal models of the world^2^. Usually, motor and reafferent signals share similar features in the spatial and temporal domains and according to the comparator model^3,4^, movements are accompanied by prediction signals (of their sensory consequences), which are compared with the actual sensory feedback in a feed-forward manner. Under such conditions, spatiotemporal congruence between predicted and reafferent sensory signals is generally associated with self-attribution of the action^5,6^ and the sense of agency: the feeling of being in control of one’s movement^5,7^. A wealth of data has shown that incongruences or sensorimotor conflicts between predicted and reafferent sensory signals lead to the loss of agency and control^8–14^.

Sensorimotor conflicts are also known to alter the perception of accompanying sensory signals. Processing of self-generated stimuli is known to be attenuated and proposed to result from a prediction-based cancelation of reafferent sensory signals^4,15,16^. A well-known example is the sensory attenuation of self-generated touch: touches produced by oneself are perceived as weaker compared to externally produced ones, even if applied with the same intensity^17–19^. Moreover, sensorimotor conflicts accompanying self-generated touches can abolish self-attenuation and thus alter the associated tactile perceptions ^4,20,21^.

Perceptual alterations caused by sensorimotor conflicts of upper-limb movements have also been observed in sensory domains other than somatosensation. For instance, studies reported a change in loudness perception of self-generated tones (by a button press), compared to tones presented passively^22–25^, which was associated with attenuated neural responses^26–30^. Recent studies have demonstrated that such auditory-motor self-attenuation effects can also be obtained for more complex sounds, such as voices^31,32^. Together, these studies show that motor activity (e.g. a button press) causally associated with the auditory feedback (e.g. a beep or the sound of one’s voice) can cause perceptual alterations of the latter through a manipulation of its spatiotemporal contingencies.

In general, most of the previous work on sensory alterations based on sensorimotor processes has focused on the investigation of sensory cues for upper-limb actions (e.g. pressing a button). However, the concept of agency, sensorimotor processes and the comparator model have also been applied to movements of the body as a whole (e.g. gait)^33–35^, thus affecting the full-body sensorimotor system associated with self-consciousness^36,37^. Extending previous robotic designs^17,18,20^, Hara et al.^38^ associated upper-limb sensory prediction signals with reafferent sensory signals at the participants’ torso in order to alter the representation of this global, torso-centered bodily system. Using this robotic device, Blanke et al.^39^ were able to induce in healthy volunteers systematic changes in illusory own body perceptions (i.e. self-touch) and mild psychosis-like phenomena that depended on sensorimotor conflicts. Specifically, while applying conflicting sensorimotor stimulation between upper-limb movements and tactile feedback on the back participants reported stronger somatic passivity (i.e. that tactile sensations are being imposed upon their body by somebody else) and felt being in a presence of a non-existing alien entity, phenomenologically resembling passivity experiences^40–42^ and presence hallucinations^43–45^ observed in schizophrenia.

Here, we investigated whether such robotically-mediated sensorimotor conflicts that are able to induce a mild psychosis-like state^39^ can also alter voice perception. Alterations of voice perception are highly prevalent in schizophrenia in the form of auditory verbal hallucinations (AVH) – i.e. hearing voices in the absence of a speaker. Given the importance of the comparator model both for somatic passivity and AVH^46–48^, we wanted to explore whether robotically-mediated sensorimotor conflicts in healthy participants induce changes in voice perception, resembling the auditory alterations and experiences observed in patients with schizophrenia – specifically loudness alterations^49–51^ and self-other vocal confusion^52–54^. In two independent experiments, participants were asked to perform repeated upper-limb movements^39^, which were conveyed as tactile feedback on their back by the robotic system^38^. Participants applied sensorimotor stimulation either in a synchronous manner or with a delay while they also performed either the loudness or the self-other voice discrimination task.

## Methods

### Participants

Each of the two separate experiments involved 30 healthy participants from the general population. In experiment 1, 9 participants were male (mean age ±SD: 21.8±2.4 years) and in experiment 2 14 participants were male (23.7±2.4 years). All participants were right-handed according to Edinburgh Handedness Inventory, fluent in French, and without any hearing deficits. Before participating in the experiment, they were screened for eligibility criteria by means of an anamnestic interview investigating medication and substance use, as well as a personal and family history of psychiatric or neurological disorders. Participants were naive to the purpose of the study, gave informed consent in accordance with institutional guidelines (Research project approved by the Comité Cantonal d’Ethique de la Recherche of Geneva) and the Declaration of Helsinki, and received monetary compensation (CHF 20/h).

## Procedure and materials

We conducted two experiments with the same general procedure and experimental design. Experiment 1 consisted of two and Experiment 2 of three sessions. For the first session of both experiments, participants came with an acquaintance, who also participated in the study, and their voices were recorded. For the second and third sessions (auditory tasks), participants came individually.

### Auditory tasks

Participants were recorded saying 10 words in French (see supplementary material). Audacity software was used to filter out the background noise and to normalize the recordings for average intensity (−12 dBFS) and duration (500 milliseconds). The pre-processed voice recordings were then entered into TANDEM-STRAIGHT^55^ to generate voice morphs between two participants (e.g. a voice morph could contain 40% of person A’s, 60% of person B’s voice). Finally, copies of the voice morphs with different sound intensities were created and the resulting audio files were played to participants through a JBL Control 1 Pro speaker placed 1 meter behind them.

During both auditory tasks (loudness, self-other), blindfolded participants repeatedly heard the same word twice, separated by 500 milliseconds. In the loudness task, both words contained the same ratio of the two voices (50% of both participants), but differed in sound intensity. In the self-other task, both words were equally loud, but contained a different ratio of the two voices. In the loudness task, participants reported which of the words they perceived as louder and in the self-other task which of the two words sounded more like their own voice.

Unbeknown to the participants, the first word in each word-pair always sounded the same (50% self-voice, -12 dBFS). The second word varied, either in sound intensity (for the loudness task) or in self-voice percentage (for the self-other task). Six sound intensity levels (dBFS: -14, -13, - 12.5, -11.5, -11, -10) and six voice ratios (% self-voice: 15, 30, 45, 55, 70, 85) were chosen based on extensive pilot testing.

### Robotic system

The robotic system consisted of two integrated units: the front part – a commercial haptic interface (Phantom Omni, SensAble Technologies) – and the back part – a three degree-of-freedom robot^38^. Participants were seated between the front and back robot and were asked to perform repeated poking movements with their right index finger using the front robot, which was replicated by the movements of the back robot, which applied corresponding touches on their back. This was done either in synchronous (without delay) or asynchronous (with 500 milliseconds delay) fashion, creating different degrees of sensorimotor conflict between the upper limb movement and somatosensory feedback on the back^39^.

Experiment 1 and consisted of synchronous and asynchronous sensorimotor conditions. Experiment 2 contained two additional conditions. In the motor-baseline condition, participants performed movements on the front unit, but did not receive the corresponding somatosensory feedback by the back unit. In the touch-baseline condition, the experimenter (not the participant) performed the movements on the front unit, but the participant received the corresponding somatosensory feedback by the back unit. These two conditions served as baselines, as there was no sensorimotor coupling.

In experiment 2 we also tested whether torso-centered tactile feedback (i.e. back) was necessary for the present effects^37^. For this, we added two more conditions in which the same setup was used as in the synchronous and asynchronous conditions, except that tactile feedback was not applied to the back but to the left hand of the participants – i.e. the back unit was placed in front of the participants and adjusted to point downwards in the vertical axis in order to touch their left hand.

### Experimental design

In experiment 1, participants performed two blocks of each auditory task (loudness and self-other) – one block in the synchronous and another block in the asynchronous condition. Each block started with 60 seconds of robot manipulation, without auditory stimulation, after which an auditory cue indicated the beginning of the actual auditory task. Throughout the auditory tasks, participants continued moving the robot. Importantly, auditory stimuli and participants’ movements were not time-locked. Each block contained 60 trials (10 words, each presented with 6 stimulus intensities) presented in a randomized order. The order of tasks (self-other/loudness) and conditions (synchronous/asynchronous) was counterbalanced across participants. An Inter-trial interval of 1 to 1.5 second (randomly jittered), was added to avoid predictability of the stimuli. (Figure 1). The experimental design was created in MATLAB 2017b with Psychtoolbox library^56–58^.

**Figure 1.**
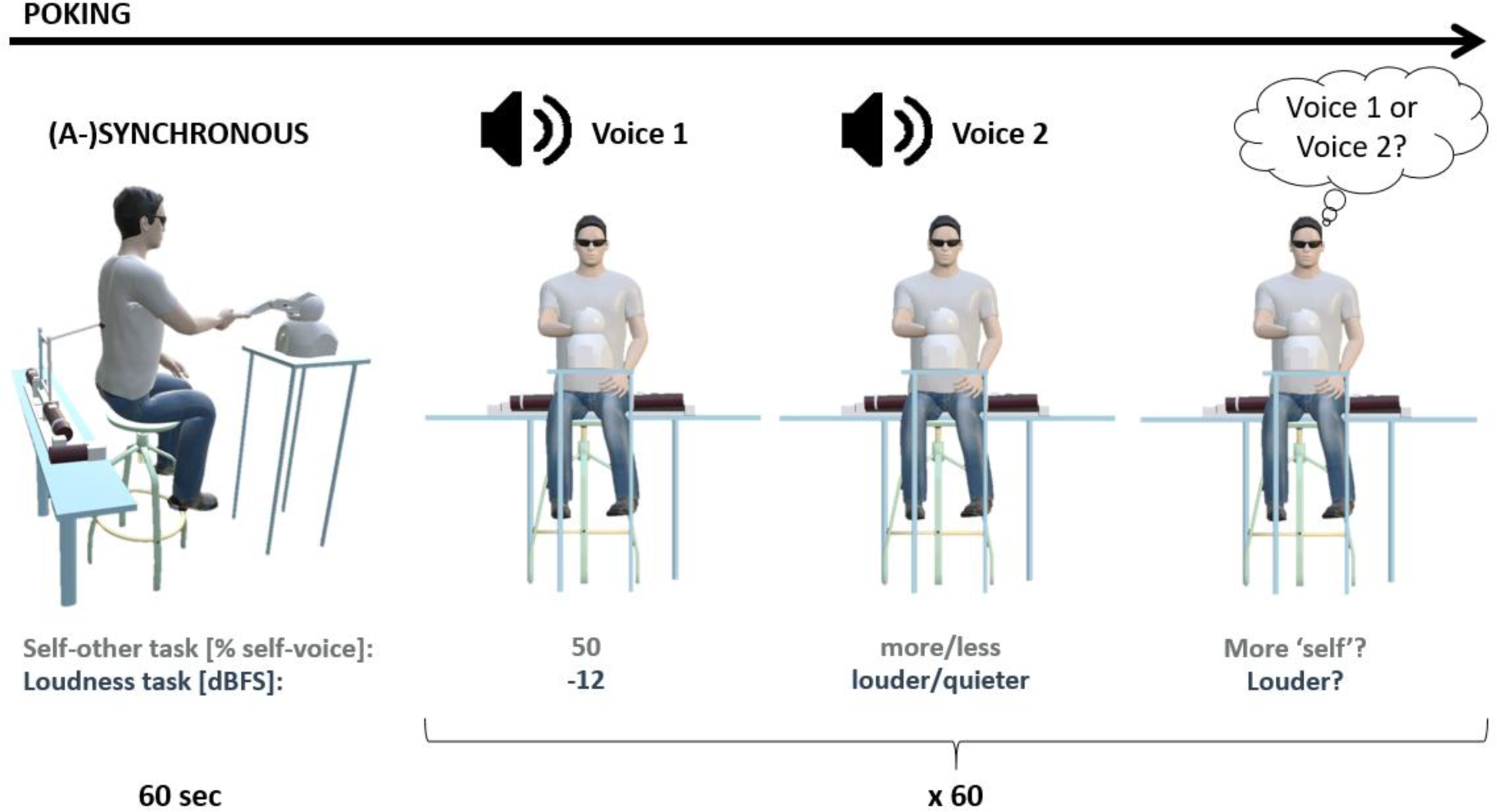
Experimental block design.

In experiment 2, participants performed four blocks of the loudness task (synchronous, asynchronous, motor-baseline, touch-baseline) and two additional hand feedback blocks (hand-synchronous, hand-asynchronous). All were equivalently designed as the loudness-task blocks of experiment 1. The order of blocks was pseudorandomized across participants. In experiment 2, there was no self-other task.

In both experiments, we additionally assessed the subjective experience evoked by the combination of robotically-mediated sensorimotor conflicts and ambiguously-voiced stimuli. Thus, after the auditory tasks, participants performed additional questionnaire blocks in which they passively listened to the same voice morphs while manipulating the robot. For each experimental condition there was an additional block after which they rated several items on a previously used questionnaire^39^ (supplementary material). In experiment 1, we added two blocks (synchronous, asynchronous). In Experiment 2 we added six blocks (synchronous, asynchronous, motor-baseline, touch-baseline, hand-synchronous, hand-asynchronous).

## Statistical analysis

Data of experiment 1 were analyzed with mixed-effects logistic regressions with Response as dependent variable and Condition (synchronous, asynchronous) and Stimulus (levels: 1-6), together with their interaction, as fixed effects. The Response-variable indicates whether participants perceived a stimulus as louder (loudness task) or as sounding more like their own voice (self-other task) compared to the reference stimulus. Random effects included a by-subject random intercept. By-subject random slopes for the main effects were added following model selection based on maximum likelihood. Trials with reaction times greater or smaller than two interquartile ranges from the median for each subject were considered as outliers and excluded.

Analysis for experiment 2 followed a similar approach (two logistic mixed-effects models with Response as a dependent variable). The first model was designed to assess the joint effects of synchrony and location of sensorimotor conflicts, including Condition (synchronous, asynchronous), Location (torso, hand) and Stimulus (levels: 1-6) with interaction terms as fixed effects. The second model extended the first one by investigating the effects of the sensorimotor coupling, regardless of the location. Therefore, it included no main effect of Location and the main effect of Condition had three instead of two levels (synchronous, asynchronous, baseline). For both experiments, a linear mixed-effects regression was also performed with Reaction Times as a dependent variable. Analysis showed no significant differences between experimental conditions (supplementary material). Questionnaire ratings were assessed by a mixed-effects linear regression and analyzed jointly for experiment 1 and 2, to increase statistical power. As fixed effects, we entered Condition (synchronous, asynchronous) and Question (q1 – q9) with interaction term into the model. As random effects, we had by-subject random intercepts. For the questionnaire items, which significantly differed between the two conditions, we conducted an additional mixed-effects linear regression investigating the fixed effect Location (torso, hand). All analyses were performed with R^59^, using notably the afex^60^, ggplot2^61^, sjplot^62^, lme4^63^, and lmerTest^64^ packages.

## Results

### Auditory task

#### Experiment 1 (Loudness, Self-other)

A mixed-effects logistic regression on loudness judgment revealed higher intercepts in the asynchronous compared to the synchronous condition (estimate=-0.39, Z=-2.14, p=0.03). The model had a main effect of Stimulus (estimate=0.59, Z=9.50, p<0.001) and showed no interaction between the Condition and Stimulus (estimate=0.08, Z=0.05, p=0.12). To further investigate the Stimulus effect observed in the loudness task, we performed the same mixed effects logistic regression for each Stimulus level. Results showed that voices were perceived significantly louder in the asynchronous condition only for the lowest sound intensity level (estimate=-0.5, Z=-2.49, p=0.01) (Figure 2, left), whereas all other stimulus levels did not differ between conditions (supplementary material).

**Figure 2.**
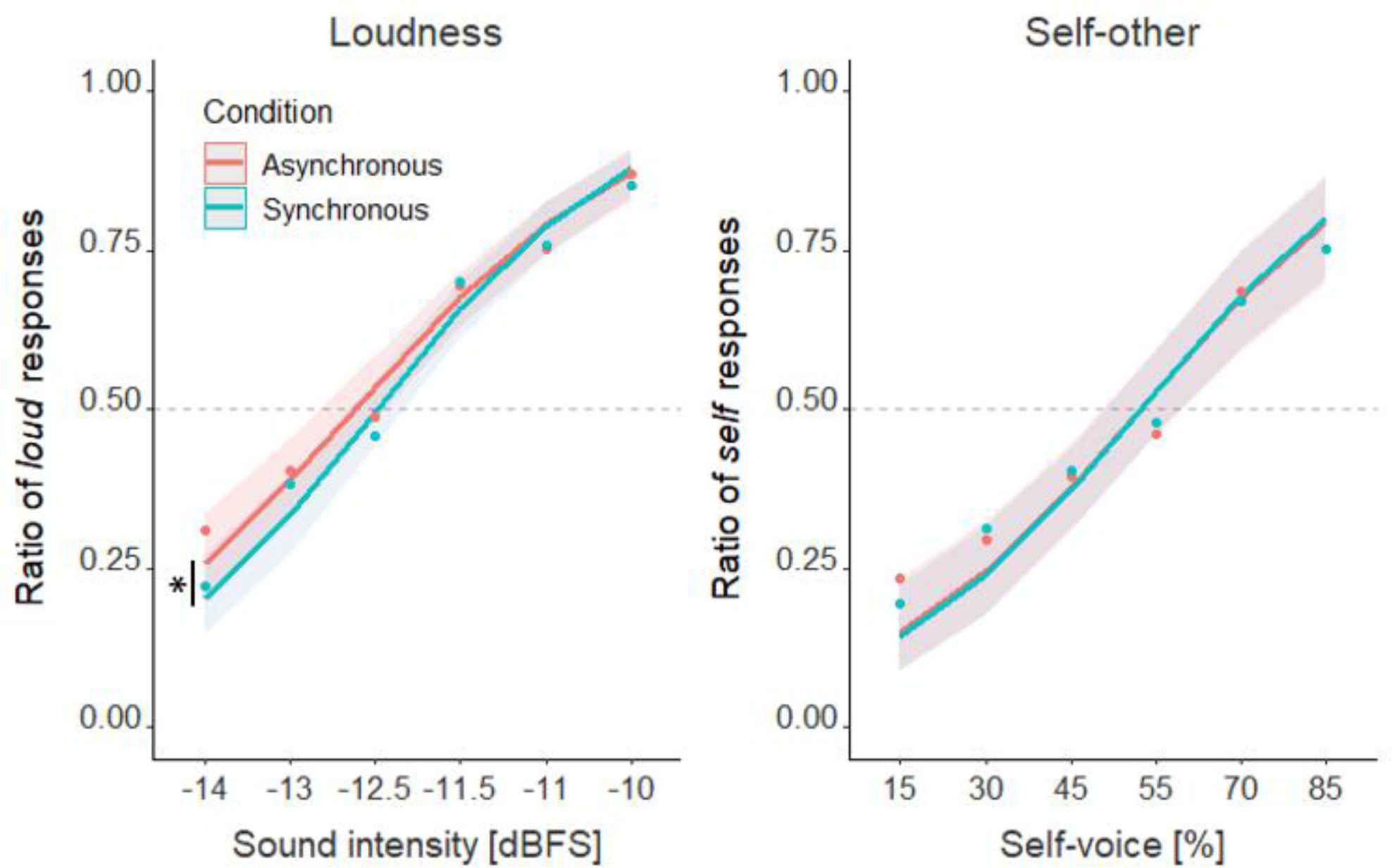
Psychometric curves fitted for the two auditory tasks of experiment 1. The points indicate the rate at which the corresponding voice was perceived as louder (Loudness task) or more resembling own voice (Self-other task) than the baseline. The shaded areas around each curve represent the 95% confidence intervals. Intercept was significantly higher in the asynchronous condition and for the loudness task only, indicating that the quieter voices were perceived as louder. *: p<0.05.

Concerning the self-other discrimination task, a mixed-effects logistic regression indicated a main effect of Stimulus (estimate=-2.36, Z=-6.46, p<0.001). Intercepts of the synchronous and asynchronous conditions did not differ in the self-other task (estimate=-0.07, Z=-0.36, p=0.72), nor was there a significant interaction between the Condition and Stimulus (estimate=0.02, Z=0.36, p=0.72).

#### Experiment 2 (Loudness, hand vs torso)

Experiment 2 replicated the loudness effect observed in experiment 1. In the model assessing both the synchrony and location of sensorimotor conflicts, the intercepts were again significantly higher in the asynchronous compared to synchronous condition (estimate=-0.49, Z=-2.92, p<0.01). The responses differed across stimuli (estimate=0.36, Z=11.22, p<0.001), but there was no significant effect of Location (hand vs. torso) (estimate=-0.3, Z=-1.65, p=0.1). We observed a significant interaction only between the effects of Condition and Stimulus (estimate=0.12, Z=2.51, p=0.01). Interactions between Condition and Location (estimate=0.37, Z=1.54, p=0.12), Stimulus and Location (estimate=0.03, Z=0.69, p=0.49) and a three-way interaction between Condition, Location and Stimulus (estimate=-0.07, Z=-1.11, p=0.27) were not significant.

Analogously to experiment 1, we performed the same mixed effects logistic regression for each Stimulus level, confirming that the difference in loudness perception between the conditions occurred only for the lowest sound intensity level (estimate=-0.35, Z=-2.66, p<0.01, for other levels see supplementary material).

We next addressed the effects of the sensorimotor stimulation, regardless of feedback location. In this model the intercept in the asynchronous condition was higher than the synchronous (estimate=-0.29, Z=-2.23, p=0.03) and the baseline (estimate=-0.51, Z=-3.34, p<0.001), whereas there was no difference between the synchronous and baseline conditions (estimate=-0.17, Z=-1.29, p=0.2) (Figure 3).

**Figure 3.**
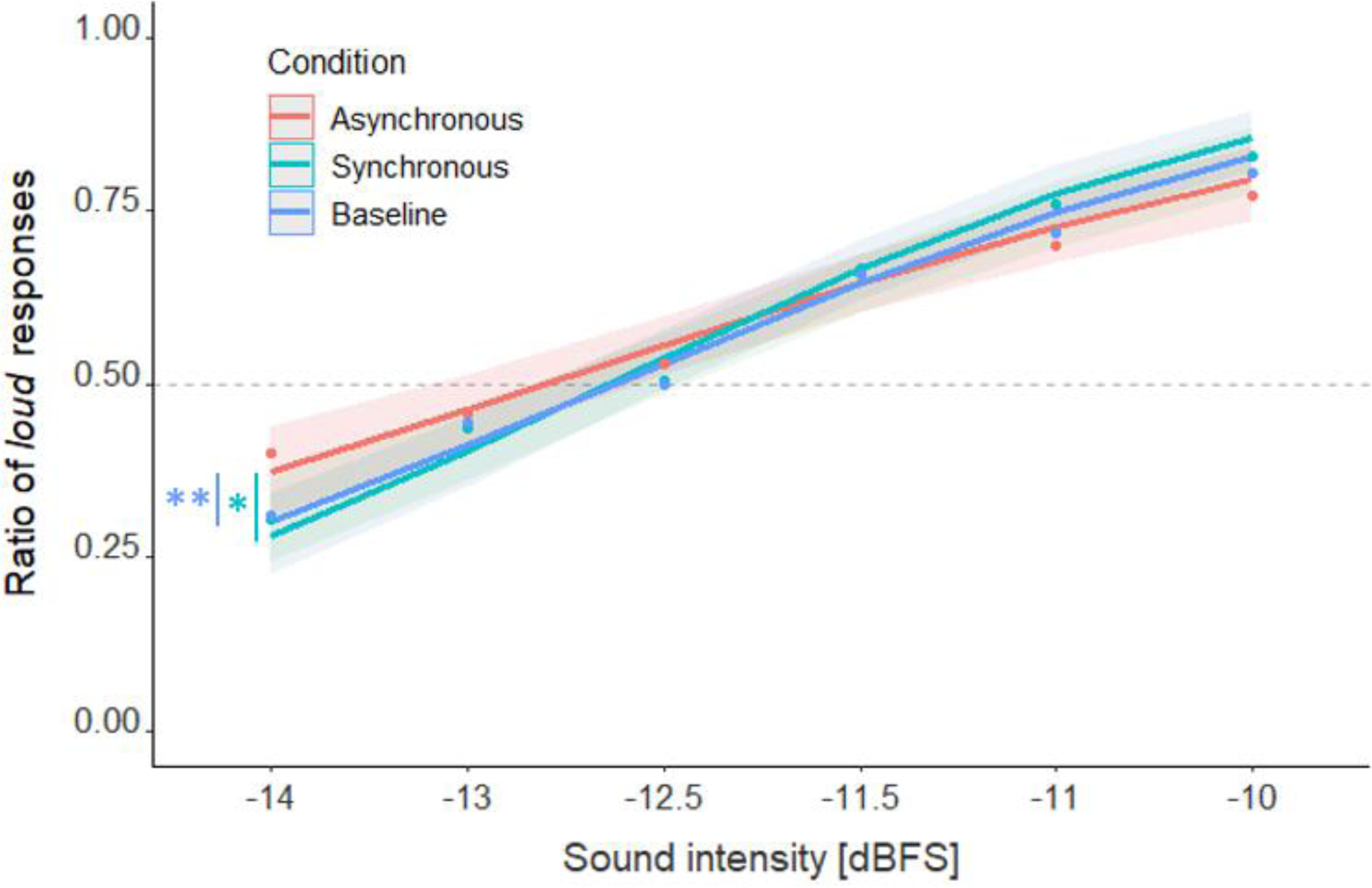
In experiment 2, intercept in the asynchronous condition was significantly higher than in the synchronous and the baseline conditions, whereas there was no difference between the synchronous and the baseline conditions. *: p<0.05, **: p<0.01.

### Subjective experience

The linear mixed-model analysis revealed that participants experienced stronger somatic passivity in the asynchronous versus synchronous condition (Figure 4A) (estimate=-0.83, t(66.94)=-2.88, p<0.01) and rated illusory self-touch significantly stronger in the synchronous versus asynchronous condition (Figure 4B) (estimate=0.64, t(67.54)=2.56, p=0.01), without any significant difference between conditions in other questionnaire items (all p>0.05).

**Figure 4.**
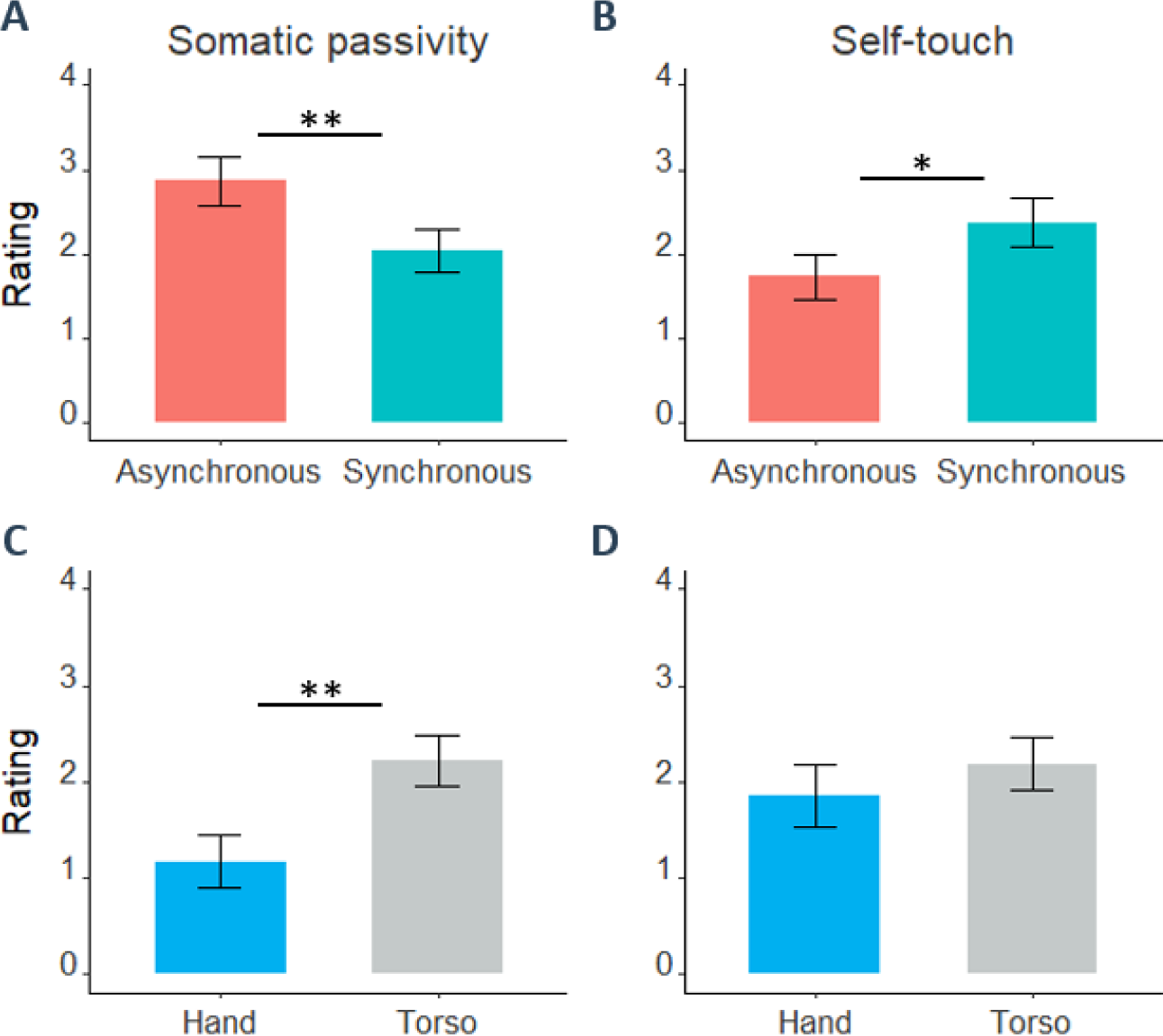
Abscissa of every bar plot indicates either the two experimental conditions (A, B: Synchronous, Asynchronous) or the location of sensorimotor conflicts (C, D: Hand, Torso) and ordinate the corresponding Likert-scale ratings. Height of a bar plot indicates the mean rating and error bars its standard error. Somatic passivity sensations were reported significantly higher in the asynchronous condition (**A**) and with sensorimotor conflicts applied to torso compared to hand (**C**). Self-touch impressions were stronger in the synchronous condition (**B**) but equally strong for both locations (**D**). *: p<0.05, **: p<0.01

For the two significant questionnaire items, an additional mixed-effects linear regression was applied, in order to investigate the effects of Location, showing that somatic passivity was significantly stronger when sensorimotor conflicts were applied on the torso vs. hand (Figure 4C) (estimate=1.34, t(88.56)=3.08, p<0.01). Self-touch ratings did not differ between the two locations (Figure 4D) (estimate=-0.1, t(87.66)=-0.24, p=0.81).

To assess the relationship between subjective experience and auditory perception, we ran mixed-effects logistic regression with significant questionnaire items (Passivity and Self-touch) as additional factors and divided participants in two groups – those with a positive asynchronous-synchronous rating difference (Passivity+, Self-touch+) and those with a negative or zero difference (Passivity-, Self-touch-). Model showed a significant interaction between Passivity and Condition (estimate=0.39, Z=2.04, p=0.04) (supplementary material). Investigation of the interaction showed that loudness perception was altered only in Passivity-group (Figure 5, left) (Condition: estimate=-0.54, Z=-3.71, p<0.001; Stimulus: estimate=0.47, Z=7.07, p<0.001; Condition-Stimulus interaction: estimate=0.12, Z=3.05, p<0.01), with no difference between conditions in Passivity+ group (Figure 5, right) (supplementary material). There were no significant interactions between Self-touch and Condition (supplementary material).

**Figure 5.**
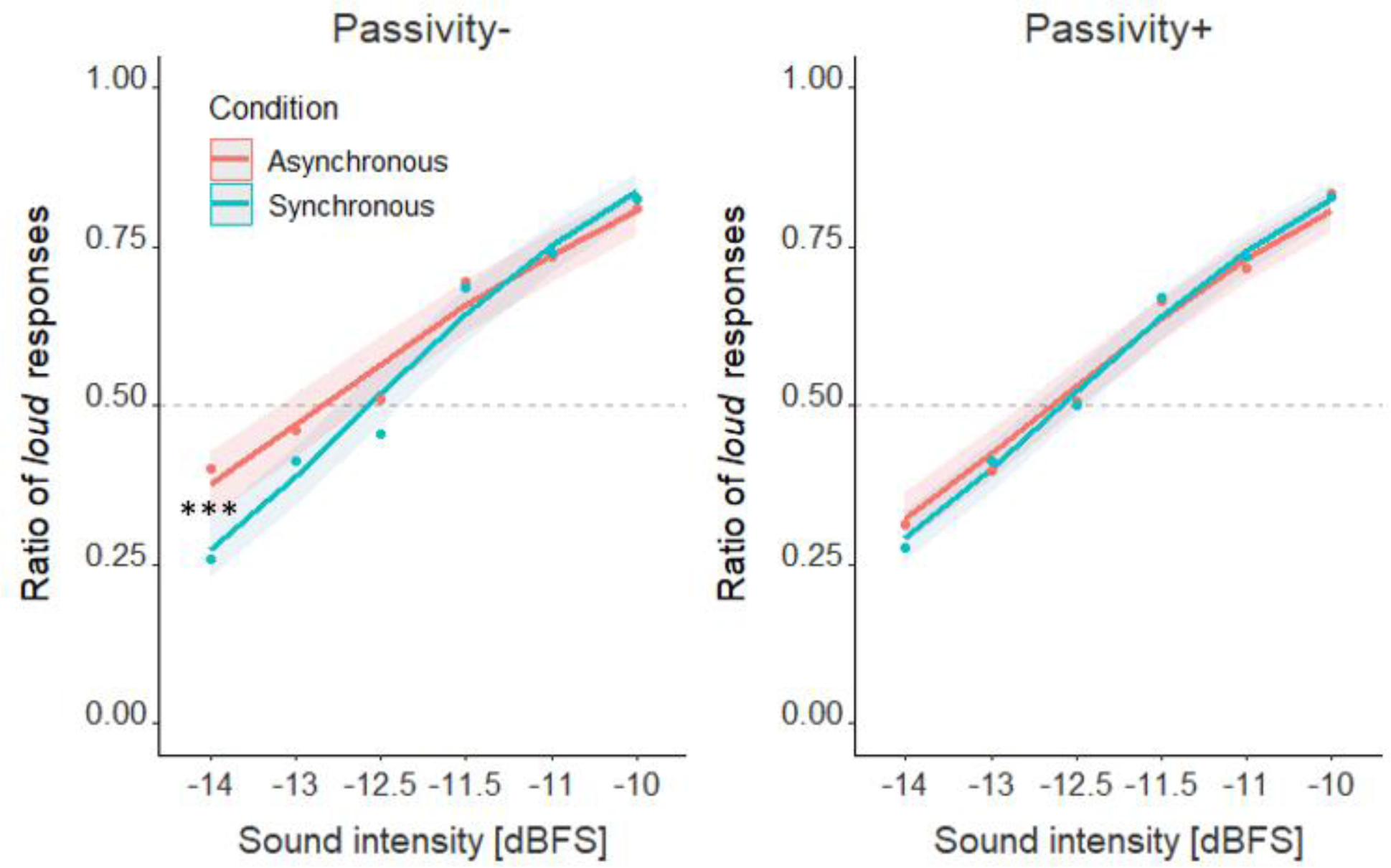
Quiet voices were amplified only for the participants not experiencing somatic passivity during the experiment (Passivity-, left). With somatic passivity (Passivity+, right) there was no change in voice perception. ***: p<0.001

## Discussion

Replicating the induction of somatic passivity based on sensorimotor stimulation in a healthy population using a robotic procedure^39,65,66^ we investigated potential links with voice perception and clinical phenomenology (i.e. AVHs) and demonstrate that voice perception is modulated by sensorimotor stimulation with somatosensory feedback. We confirmed this somatosensory-motor effect on auditory perception in two independent cohorts in two studies. Specifically, quiet voices were perceived as louder in the asynchronous condition, differing from voices heard in synchronous and baseline conditions.

Changes in perception during actions are usually interpreted within the comparator model framework: self-generated movements are accompanied by sensory predictions, which cause an attenuation of the reafferent sensory signals, especially if they are received in spatiotemporal congruency^3,4^. Thus, in order for the sensory signal to be attenuated, predicted and reafferent sensory consequences must have a consistent spatiotemporal relationship such as pushing a response button with one’s right index finger attenuating processing of tactile stimuli at the fingertip, via predictive sensory signaling^4,17^. Related work has extended these findings to auditory perception, showing that auditory processing of a sound triggered by a button press is attenuated^27,31^. Lack of predictive mechanisms is associated with decreases in sensory attenuation and perceived as amplification of the sensory stimuli accompanying actions (stronger touches^17,67,68^; louder sounds^22–25^).

The present findings extend sensory attenuation research in two ways. First, there was no time-locking between our participants’ movements and the auditory stimuli they were asked to judge. Participants manipulated the robot independently from the sounds and the auditory task – ruling out the possibility that classical effects linked to the comparator framework and associated with trial-by-trial sensory comparisons between an action and its sensory consequences account for the present effects. Secondly, perceptual changes in both experiments were only present in the asynchronous condition, accentuating the importance of temporal aspects (between movement and somatosensory feedback) of sensorimotor conflicts. In experiment 1 and 2, we observed a difference in loudness perception between the asynchronous and synchronous conditions and in experiment 2, additionally, observed that perception in the asynchronous condition is the deviating one, as it alone differed from baseline conditions. Crucially, the perception in the spatially-conflicting, yet synchronous condition did not differ from the no-conflict conditions (touch- and motor-baseline), suggesting that mainly the temporal conflict, present only in the asynchronous condition, drives the present perceptual effects. Temporal conflicts have been shown to cause a loss of agency, by manipulating sensory action consequences of upper-limb movements and related losses of hand movement agency^8–14^. When extending such manipulations to a torso-centered bodily system^36,37^, other-agency changes can be introduced^39,66^, together with a state of an altered bodily self-consciousness, including the alien agent^39,66^. We argue that loudness amplification, observed solely in the asynchronous condition, may represent a reduction of auditory self-attenuation, resulting from such other-agency-related alterations in bodily self-consciousness.

Deficits in self-attenuation have been observed in schizophrenia. When healthy participants overestimate the externally-applied stimulation, arguably due to sensory attenuation for actively produced movements, individuals with schizophrenia perform differently, suggesting an alteration of corrections related to sensory attenuation^69,70^, compatible with neural responses between self- and externally-generated sounds in individuals with schizophrenia^46,47^ and in healthy individuals depending on hallucination proneness^67,71,72^. Our results in healthy subjects support this inverse relationship by demonstrating a lack of loudness increase only in the hallucinating group, extending previous data on changes in self-other voice discrimination in early psychosis patients with passivity symptoms^66^. Interestingly, somatic passivity was experienced more strongly when receiving torso-centered bodily feedback, compared to hand feedback, a finding not observed for illusory self-touch. As, in addition, the strength of illusory self-touch did not interact with the loudening effect, we suggest that torso-centered manipulations involving sensations related to another agent (passivity experience) interfere more strongly with voice perception than more focal somatosensory feedback (hand). Collectively, these findings suggest that asynchronous torso-centered sensorimotor stimulation (1) induces a mild psychosis-like state in the form of somatic passivity and (2) is associated with a loudening of voices, however, that (3) experiencing somatic passivity leads to a lack of voice loudening, suggesting a reduction in self-attenuation mechanisms.

Differences in divided attention between asynchronous vs. synchronous conditions cannot account for these effects, because (1) both sensorimotor conditions contained a strong conflict and both induced an altered mental state (asynchronous: somatic passivity; synchronous: self-touch), because (2) reaction times revealed no differences between both sensorimotor conditions, and because (3) the effect was only observed in one auditory task. Although it is further known that auditory perception is altered during movement^73^, movements in the synchronous and motor-only conditions were not accompanied by changes in auditory perception, suggesting the necessity of a temporal conflict for the present loudness effect. The present sensorimotor conflicts did not affect self-other voice discriminability. It is possible that a motor component involving speech production is necessary to observe a misattribution of one’s own voice in healthy individuals, as is argued to occur in AVHs^52,74–76^. The orthogonal sensorimotor stimulation, as tested in the present experiments, changes loudness, but not identity of the heard voice.

To the best of our knowledge, this is the first study to demonstrate that temporal sensorimotor conflicts in the somatosensory domain can affect voice perception even if the auditory stimulus is not systematically linked to the movement. We found that healthy listeners heard quiet voices as louder when exposed to asynchronous sensorimotor stimulation related to somatic passivity experiences. We argue that this amplification represents a reduction in self-attenuation mechanisms, reminiscent of altered voice perception in psychiatric populations. Together, our findings extend the understanding of subjective and perceptual alterations caused by conflicting sensorimotor processing and suggest that passivity experiences and voice perception rely, at least partly, on common sensorimotor brain mechanisms.

## Author Contributions

Study concept and design: PO, GR, NF, OB. Acquisition of data: PO. Analysis and interpretation of data: PO, GR, NF, OK, OB. Drafting of the manuscript: PO, OK, NF, OB. Critical revision of the manuscript for important intellectual content: All authors.

Statistical analysis: PO, GR, NF. Obtained funding: OB. Administrative, technical, or material support: All authors. Study supervision: NF, OB.

