## Supplementary material for "Sensorimotor conflicts induce somatic passivity and louden quiet voices in healthy listeners"

Running title: Sensorimotor basis of voice perception

**Pavo Orepic[1], Giulio Rognini[1], Oliver Alan Kannape[1], Nathan Faivre[2,*], Olaf Blanke[1, 3, *]**

[1] Laboratory of Cognitive Neuroscience, Center for Neuroprosthetics and Brain Mind Institute, Faculty of Life Sciences, Swiss Federal Institute of Technology (EPFL), Switzerland

[2] Laboratoire de Psychologie et Neurocognition (LPNC), CNRS UMR 5105, Université Grenoble Alpes, France

[3] Faculty of Medicine, University of Geneva, Geneva, Switzerland

* equal contribution

**Corresponding author:**

Olaf Blanke

Bertarelli Chair in Cognitive Neuroprosthetics

Center for Neuroprosthetics & Brain Mind Institute

School of Life Sciences

Campus Biotech

Swiss Federal Institute of Technology

Ecole Polytechnique Fédérale de Lausanne (EPFL)

CH – 1012 Geneva

### Words

Participants were recorded saying 10 words in French (*clou, fouet, hache, lame, lutte, os, rat, sang, scie, ver*). The words were chosen from the list of 100 negatively-valenced words, as rated by 20 schizophrenic patients and 97 healthy participants[1].

### Experiment 1 (Loudness, Self-other)

In the loudness task, participants perceived the target voice as louder than the reference in 57.4% of all trials, with an average of 26.6% for the lowest (-14 dBFS) and 86.2% for the highest stimulus level (-10 dBFS), indicating that we effectively sampled the parameter space of the task (main Figure 2, left). In the self-other task, participants perceived their voice as the dominant one in 46.9% of trials, with an average of 21.4% for the lowest (15% self-voice present) and 75.1% for the highest stimulus level (85% self-voice) (main Figure 2, right).

To further investigate the effect observed in the loudness task, we performed the same mixed effects logistic regression for each Stimulus level, allowing us to identify the sound intensity levels driving the difference in loudness perception. Results showed that voices were perceived significantly louder in the asynchronous condition only for the lowest sound intensity level (quiet sounds; level 1: estimate=-0.5, Z=-2.49, p=0.01), whereas all other stimulus levels did not differ between conditions (2: estimate=-0.11, Z=-0.62, p=0.53; 3: estimate =-0.13, Z =-0.79, p =0.43; 4: estimate=0, Z=0.03, p=0.98; 5: estimate=0.04, Z=0.19, p=0.85; 6: estimate=-0.15, Z=-0.6, p=0.55 ). Thus, quiet voices were perceived as louder in the asynchronous condition, whereas there were no perceptual differences for louder voices between the two experimental conditions (main Figure 2, left).

### Experiment 2 (Loudness, hand vs torso)

Participants perceived the target voice as louder than the reference in 57.9% of all trials, with an average of 34.9% for the lowest (-14 dBFS) and 79.5% for the highest stimulus level (-10 dBFS) (main Figure 3).

Analogously to experiment 1, we performed the same mixed effects logistic regression for each Stimulus level, confirming that the difference in loudness perception between the conditions occurred only for the lowest sound intensity level (quiet sounds; level 1: estimate=-0.35, Z=-2.66, p<0.01). All other stimulus levels did not differ between both sensorimotor conditions (2: estimate=-0.02, Z=-0.18, p=0.86; 3: estimate=-0.06, Z=-0.53, p=0.6; 4: estimate=-0.03, Z=-0.2, p=0.84; 5: estimate=0.11, Z=0.13, p=0.4; 6: estimate=0.14, Z=0.92, p=0.36).

The model investigating the effects of the sensorimotor coupling, regardless of the location, could be designed in one more way, as it is mentioned in the main text. Namely, the main effect of Condition contained four instead of three levels (synchronous, asynchronous, motor-baseline, touch-baseline). In this model the intercept in the asynchronous condition was significantly higher than the ones from all other conditions (synchronous: estimate=-0.31, Z=-2.56, p=0.01; motor-baseline: estimate=-0.4, Z=-2.42, p=0.02; touch-baseline: estimate=-0.59, Z=-3.57, p<0.001). There were no other significant differences between the intercepts of the other three conditions (synchronous and motor-baseline: estimate=-0.09, Z=-0.52, p=0.61; synchronous and touch-baseline: estimate=-0.28, Z=-1.68, p=0.09; motor-baseline and touch-baseline: estimate=-0.2, Z=-0.1, p=0.32). Thus, this model confirms the findings of the model mentioned in the main text.

### Reaction times

For both experiments, a linear mixed-effects regression was also performed with Reaction Times as a dependent variable. The model contained the same fixed and random effects as the model with Response as a dependent variable, except that a polynomial expansion of Stimulus variable to the power of two was added, to account for the non nonlinear distribution of reaction times around the point of subjective equivalence (i.e., reaction times became shorter with more extreme stimulus levels).

Participants responded to the auditory stimuli on average in 1.23 seconds in the loudness task and in the self-other task in 1.42 seconds. In experiment 1, there were no significant differences in reaction times between the conditions in any of the tasks (loudness: estimate = 0, $t(30) = 0.26$, $p = 0.8$; self-other: estimate = 0, $t(29.7) = -0.12$, $p = 0.9$). There was a main effect of Stimulus in both tasks (loudness: estimate = -3.21, $t(3462) = -5.86$, $p < 0.001$; self-other: estimate = -3.36, $t(3414) = -5.41$, $p < 0.001$), without significantly interacting with the conditions (loudness: estimate = 0.96, $t(3462) = 1.29$, $p = 0.2$; self-other: estimate = -1.34, $t(3414) = -1.53$, $p = 0.13$).

In experiment 2, there were no significant effects of Condition (estimate = -0.02, $t(46) = -0.63$, $p = 0.53$) nor Location (estimate = 0.05, $t(36) = 1.29$, $p = 0.21$) on reaction times. The model showed a main effect of Stimulus (estimate = -2.27, $t(6818) = -2.48$, $p = 0.01$), but no significant interactions between the fixed effects (Condition and Location: estimate = -0.02, $t(6822) = -0.87$, $p = 0.39$; Condition and Stimulus: estimate = 0.38, $t(6818) = 0.29$, $p = 0.77$; Location and Stimulus: estimate = 0.21, $t(6818) = 0.17$, $p = 0.87$; Location, Condition and Stimulus: estimate = 0.67, $t(6817) = 0.37$, $p = 0.71$). As in experiment 1, reaction times did not differ between conditions.

### Questionnaire

After each questionnaire block, participants filled in the following 9-item questionnaire (extended from Blanke et al., 2014[2]) by rating how strongly they agreed with each item on a Likert scale from 0 (Not at all) to 6 (Very strong). Participants reported the subjective feeling of touching oneself (“I felt as if I was touching my back by myself.”), the strength of somatic passivity (“I felt as if someone else was touching my back.”) and presence hallucination (“I felt as if someone was standing close to me.”). Additionally, we investigated whether sensorimotor conflicts and voiced stimuli affected each other on a subjective level. On one hand, we examined whether the voiced stimulation imposed an identity to the potentially evoked presence (“I felt as if my friend was standing close to me.”; “I felt as if someone else than my friend was standing close to me.”). On the other hand, we explored whether sensorimotor conflicts biased the perceived auditory ambiguity (“I felt as if I heard my friend’s voice more often than my own voice.”; “I felt as if I heard a voice that was neither my friend’s nor mine.”). Finally, participants were asked to report changes in bodily sensations experienced during the corresponding block (“While hearing the words, I felt changes in my body sensations (e.g. lighter, warmer, I felt tingling sensations etc.)”), and to describe them by means of an open self-report (“If answer is between 1 and 6, please describe the changes in body sensations.”).

### Spontaneous negative reports

As a final part of our questionnaire, there was an open-type question, where participants were asked to describe the bodily sensations they might have experienced during the experiment. Several participants freely reported about emotional changes and 11 participants felt negative emotions and, in addition, reported them only during the asynchronous condition (I felt "annoyed", "anxious" (2), "bad", "choking", "painful", "sad", "stressed" (2), "tense", "worried") (only one instance was reported during the synchronous condition ("bad")). A chi-square test of independence was performed to examine the relation between experimental conditions (synchronous, asynchronous) and emotional valence (negative, non-negative). This test revealed that negative emotions were more likely during the asynchronous condition ($X^2$ (2, N=120) = 9.25 p < .01).

Out of 60 participants, 31 (52%) had higher somatic passivity in the asynchronous condition (Passivity+ group, main Figure 5, right) and all participants who spontaneously reported negative emotions in the asynchronous condition also belonged to the Passivity+ group. 23 (38%) participants reported higher self-touch sensations in synchronous condition.

Thus, the robot-induced sound effect and psychosis-like state were associated with negative emotional valence, as participants spontaneously reported negative emotions (Supplemental Figure 1), especially during the asynchronous condition and in individuals experiencing somatic

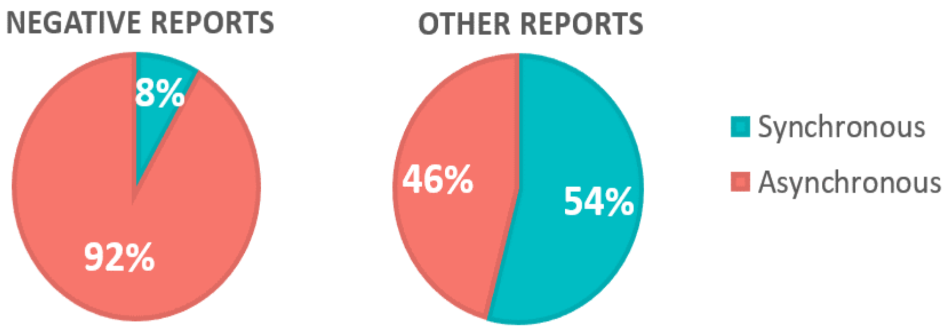


**Supplemental Figure 1**. Open self-reports. When asked to describe changes in bodily sensations – such as feeling warmer or lighter – experienced during the experimental blocks, some participants spontaneously reported feeling negative emotions. Such negative reports were associated with the asynchronous condition.

passivity (Passivity+). One could argue that the appearance of negative emotions was due to the fact that the participants were hearing negative words, yet all experimental conditions contained the same negative auditory stimuli, and negative sensations were reported predominately after the asynchronous condition. This represents another phenomenological resemblance to clinical voice-hearing, as negative valence has been proposed to be a determining factor for separating clinical from healthy AVHs[3–5].

### Auditory perception and subjective experience

To assess the relationship between the subjective experience and auditory perception, we ran the same mixed-effects logistic regression with significant questionnaire items (Passivity and Self-touch) as additional factors. For both items, participants were divided in two groups – those with a positive difference in ratings between the asynchronous and synchronous conditions (Passivity+ and Self-touch+) and those with a negative or zero difference (Passivity- and Self-touch-).

The model which had Passivity as an additional factor showed significant effects of Condition (estimate=0.59, Z=-4.27, $p<0.001$) and Stimulus (estimate=0.39, Z=14.88, $p<0.001$), with a significant interaction between the two factors (estimate=0.13, Z=3.56, $p<0.001$). The effect of Passivity borderlined with significance (estimate=-0.28, Z=-1.65, p=0.09) and interacted with the effect of Condition (estimate=0.39, Z=2.04, p=0.04). Passivity did not interact with Stimulus (estimate=0.05, Z=1.37, p=0.17) and the three-way interaction between Condition, Stimulus and Passivity was not significant (estimate=-0.08, Z=-1.54, p=0.12).

Investigation of the interaction between Passivity and Condition showed that loudness perception was altered only in Passivity- group (main Figure 5, left) (Condition: estimate=-0.54, Z=-3.71, p<0.001; Stimulus: estimate=0.47, Z=7.07, p<0.001; Condition-Stimulus interaction: estimate=0.12, Z=3.05, p<0.01), with no difference between conditions in Passivity+ group (Figure 5, right) (the effect of Condition: estimate=-0.15, Z=-1.05, p=0.29; Stimulus: estimate = 0.51, Z = 9.57, p < 0.001; Condition and Stimulus interaction: estimate = 0.04, Z = 1.17, p = 0.24).

The model with Self-touch as an additional factor also showed significant effects of Condition (estimate=-0.43, Z=-3.52, p<0.001) and Stimulus (estimate=0.41, Z=18.02, p<0.001), also with a significant interaction between the two factors (estimate=0.1, Z=3.01, p<0.01). However, the effect of Self-touch was not significant (estimate=0.12, Z=0.6, p=0.55). It did not interact with the effects of Condition (estimate=0.1, Z=0.48, p=0.63) nor Stimulus (estimate=0.02, Z=0.45, p=0.65), nor was the interaction between Condition, Stimulus and Self-touch significant (estimate=-0.01, Z=-0.26, p=0.8).

Additionally, we ran monotonic (Spearman) correlation analyses between the significant questionnaire and auditory task findings. It indicated a negative monotonic relationship between the effects of synchrony on task performance and the intensity of somatic passivity (Supplemental Figure 2) ($\rho$=-0.3, p=0.03). Specifically, the difference between the asynchronous and synchronous conditions in loudness perception of quiet voices (i.e. individual average responses for the lowest stimulus level) negatively correlated with the difference in somatic passivity experienced during the asynchronous and synchronous conditions (i.e. individual questionnaire ratings). The same correlation analysis between the differences in task performance and the intensity self-touch sensation between the asynchronous and synchronous condition did not indicate a significant relationship ($\rho=0.2$, $p=0.14$).

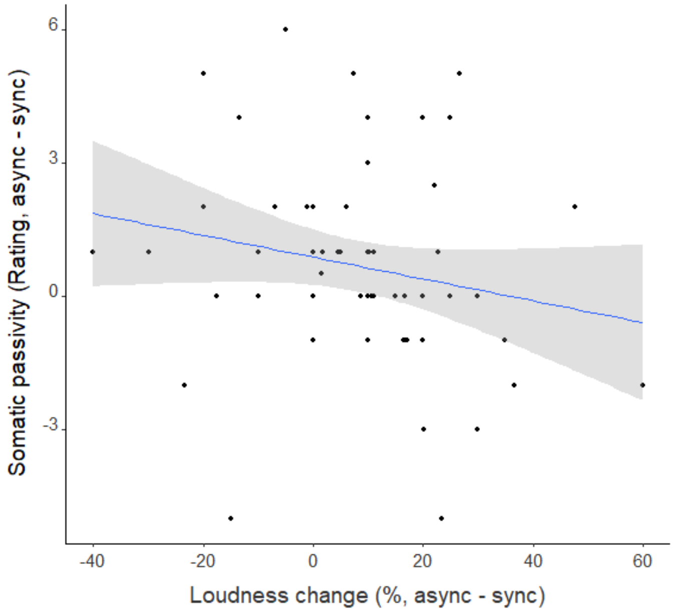


**Supplemental Figure 2**. Somatic passivity negatively correlates with voice amplification. Each dot represents an individual increase in loudness perception of quiet voices (abscissa) and somatic passivity (ordinate) between the asynchronous and synchronous conditions. Blue line represents a linear regression describing the negative monotonic relationship between the two, with the shaded area indicating its 95% confidence interval.
